# Tree species richness and evenness affect forest biomass differently across biogeographic regions

**DOI:** 10.1101/2023.12.07.570720

**Authors:** Stefania Ondei, Jessie C. Buettel, R. Zach Aandahl, Barry W. Brook, John Alroy, Luke A. Yates

## Abstract

The relationship between tree species diversity, measures of forest structure, and forest biomass has long been debated, with local- or continental-scale studies often finding contrasting results. Given the importance of forests as global carbon sinks, understanding the characteristics that underpin biomass accumulation is thus a critical component of mitigating climate change. Here we present a global analysis of 11,400 forest plots, sourced from scientific publications and forest inventories, to investigate the association of forest basal area (used as a proxy for biomass) with stem density and measures of tree species diversity (richness and evenness). We used generalised additive models to account for the confounding effects of climate and spatial signal and we modelled the density, climate, and diversity effects both globally and for each biogeographic region. Stem density showed a strong positive association with basal area across all biogeographic regions, while the effect of species richness varied. In the Palearctic, Nearctic, and Neotropical biogeographic regions, basal area was positively associated with species richness, although this was only detectable for lower values of basal area. In the Ethiopian and Oriental biogeographic regions there was no relationship between richness and basal area, while in the Australian biogeographic regions it was negative. The weak-to-no association between species evenness and basal area in all bioregions other than Australia suggests that the overall correlation emerges from processes operating at more local scales. Our results highlight the importance of accounting for biogeographic processes when evaluating strategies to mitigate climate change and support nature conservation.

## Introduction

Forests are important global carbon sinks, and, as such, a key asset in climate-change mitigation strategies. It has been estimated that forests absorb a net 7.6 billion metric tonnes of CO_2_ per year, corresponding to twice as much carbon as they emit (Harris et al., 2021). However, the potential of forests as carbon sinks is countered by ongoing land clearing, with deforestation and other land-cover changes being responsible for nearly 60% of the difference between current and potential biomass stocks (Erb et al., 2018). While reforestation practices can contribute to increase forested areas, their effectiveness in terms of biomass accumulation is difficult to assess due to the lack of understanding of the ecological drivers that maximize it. For instance, large variations in the amount of stored carbon have been detected across geographic locations (Rozendaal et al., 2022), suggesting that local environmental factors and biogeographic drivers might both affect biomass buildup (Noreika et al., 2019).

One question in forest dynamics that is yet to be resolved is whether there is a consistent association between tree density and aboveground biomass. Forest biomass is usually considered to be negatively associated with stem density, because the competition occurring as a stand develops leads to density-dependent mortality, resulting in self-thinning processes (Westoby, 1984). In line with this idea, recent research suggested that most of forest aboveground biomass is likely to be stored in its largest trees (Lutz et al., 2018, Ali et al., 2019, Cuni-Sanchez et al., 2021, Slik et al., 2013), although this effect can depend on forest types and local environmental conditions (Ku et al., 2023). However, Ligot et al. (2018) found no support for this theory in old-growth tropical forests, suggesting that local factors might also be involved. Additionally, analyses of forest structure across different environments found a positive relationship between stem density and biomass, contrary to self-thinning predictions (Schietti et al., 2016, Sagar and Singh, 2006, Kara and Keleş, 2023).

The effect of species diversity on forest-tree productivity has also long been debated. A common assumption is that the relationship between species richness and productivity is hump-shaped, with low species richness at low productivity sites because of the stress caused by harsh conditions, as well as at high productivity sites due to elevated competition (Grime, 1973). Although subsequent studies have largely confirmed this model (e.g., Potter and Woodall, 2014, Mittelbach et al., 2001), within homogeneous habitats the relationship between species richness and biomass can be positive, linear, or non-existent (Guo and Berry, 1998). A positive association between tree diversity and forest biomass, measured as either aboveground biomass, productivity, or basal area, has been found in different habitats (Huang et al., 2018, Vilà et al., 2007, Kara and Keleş, 2023, Noreika et al., 2019, Mensah et al., 2016), across large geographic areas encompassing multiple forest types (Liang et al., 2007, Vilà et al., 2013, Paquette and Messier, 2011), and in global studies (Liang et al., 2016), although the latter’s methods have been criticized (Dormann et al., 2019). Equally unresolved is the effect of species evenness, which describes the distribution of the abundance of individuals across species. Studies in plantations showed that polycultures were more productive than monospecific stands, supporting a positive association between evenness and biomass (Erskine et al., 2006, Zhang et al., 2012). However, in natural forests this association is less clear. Whilst some studies found a clear negative correlation between the two (Lee et al., 2022, Ali et al., 2018), Mulder et al. (2004) concluded that the correlation varies depending on local conditions and Fotis et al. (2018) found only a weak negative correlation.

As forest biomass is estimated to increase towards lower latitudes, as does species richness (Gillman et al., 2015), it is unclear if the potential relationship between the two is a byproduct of their co-dependence on climate. Further, the exact nature of climate’s effect on tree biomass (or productivity) is not entirely resolved. For instance, several studies have found mean annual temperature to influence tree biomass or growth, but while this effect was positive in some cases (e.g., Zell, 2018, Laubhann et al., 2009, Toledo et al., 2011), in others it was negative (Aubry-Kientz et al., 2019, Prior and Bowman, 2014, Bowman et al., 2014) or showed a temporary positive effect on tree growth, followed by a long-term negative effect D’Orangeville et al. (2018). Total annual precipitation is typically positively associated with tree biomass and growth (e.g., Slik et al., 2010, Toledo et al., 2011, Zell, 2018, Żywiec et al., 2017, Rohner et al., 2016). Climate fluctuations throughout the year have also been identified are important predictors of tree biomass and productivity (Álvarez-Dávila et al., 2017, Slik et al., 2010, Moore et al., 2006, Lebourgeois et al., 2014, Nicklen et al., 2019, Wright et al., 2018, Żywiec et al., 2017, Zhang et al., 2013, Manso et al., 2015, Toledo et al., 2011). Notably, in one broad-scale study no climate-biomass relationship was detected (Stegen et al., 2011). This combined evidence suggests that climate likely exerts a different effect on forests depending on location and the forest type investigated.

Clearly then, resolving the association of forest structure and species diversity with tree biomass is challenging due to many potential confounding effects. Global-scale analyses require data to be collected across large areas to account for climatic variations, and detailed information (e.g., basal area, species identification) must be available for each individual tree. Additionally, differences in sampling methods mean that the raw data should be accessed so as to avoid the methodological issues associated with meta-analyses (Whittaker, 2010). This was not possible until recently, when the increasing amount of research data uploaded to open-access repositories, the spread of international monitoring plot networks, and the availability of data from national forest inventories allowed for the collation of the information required (Ondei et al., 2018). Finally, the underlying statistical models must be able to adequately account for the potentially confounding effects of climate as well as the spatial location of plots, and also be flexible enough to identify different linear and non-linear effects across geographic zones.

Here we present such an analysis, based on a very large dataset of forest plots (*n* = 11,400), which covers most of the world’s forested areas and was obtained by compiling data from several national forest inventories and peer-reviewed publications. To circumvent the difficulty in calculating tree aboveground biomass for such a large number of diverse plots, which requires information on tree height and wood density, we used tree basal area as a proxy. A simple linear relationship between basal area and biomass has been found previously across different forest stands (Chiba, 1998, Qiu et al., 2021) and as such, basal area is considered a strong predictor of biomass (Mensah et al., 2016, Slik et al., 2010, Vilà et al., 2013). We used species diversity estimators to provide robust estimates of species richness and evenness and statistically accounted for the confounding effects of climate and plot location across different biogeographic regions. By properly accounting for these factors, this sought to disentangle the relationship between forest diversity and basal area from the influence of climate and geographic location.

## Methods

### Data collection and processing

Datasets were collated from the published literature, accessed independently or through The Ecological Register (http://ecoregister.org/), as well as National Forest Agencies and National Vegetation Monitoring Surveys. Each data source was individually screened to include only plots located in undisturbed primary forests (i.e., removing plantations or secondary forests) for which plot size and location were reported, as well as scientific name, number of trees and total basal area of trees with DBH ≥ 10 cm for each species present in the plot. The threshold of 10 cm was chosen as it was the most common across datasets; when a study included smaller trees those were removed before collating the data. Species names were associated with the corresponding accepted binomial name using Program R (R Core Team, 2020) and the R package *taxize* (Chamberlain and Szöcs, 2013) which relies on the Encyclopedia of Life’s Global Names Resolver (http://resolver.globalnames.org) to search and normalise names. Data from 47 scientific papers, the National Forest Agencies of seven countries (Canada, United States, Mexico, France, Italy, Sweden, and South Korea), and three National Vegetation Monitoring Surveys (Japan, New Zealand, and Australia) met our criteria. A full list of data sources and their corresponding references is provided in Supplementary Material 1.

For each plot, we calculated measures of forest diversity and structure. The number of species observed in a given plot is a function of plot size (Mittelbach et al., 2001) and this dependence is not linear (Drakare et al., 2006). To address this sampling bias, instead of accounting for plot size by incorporating it in our models, we calculated species richness using the abundance-based estimator Chao1. This provides one commonly applied approach for estimating the number of species present in the assemblage by applying a correction factor to the raw species count (Chao and Chiu, 2016, Gotelli and Colwell, 2011). As the accuracy of this estimator is sensitive to sample size, we retained only plots with at least 100 observations, or plots with a species/observation ratio equal to 10 or higher. The latter was used to ensure that plots located in typically species-poor areas (e.g., boreal forests), for which fewer than 100 observations are needed to characterize the local assemblage, were not unnecessarily excluded. Since species diversity cannot be decomposed into independent richness and evenness (Jost, 2010), we calculated Pielou’s J (Pielou, 1966) as a measure of ‘relative evenness’, defined as the evenness amount relative to the possible range given the observed richness (Jost, 2010). Total plot basal area (m^2^) and stand density, (number of trees per plot), were calculated and standardized per hectare.

Climate data was obtained from Worldclim 2.0 (Fick and Hijmans, 2017) for each plot, focusing on those variables that have been previously identified as likely drivers of basal area. This included annual precipitation (Slik et al., 2010, Toledo et al., 2011, Żywiec et al., 2017), precipitation of the active growth period (i.e., the wettest or warmest quarter; Moore et al., 2006, Nicklen et al., 2019), precipitation seasonality (Slik et al., 2010, Álvarez-Dávila et al., 2017), mean annual temperature (Álvarez-Dávila et al., 2017, Aubry-Kientz et al., 2019, Zell, 2018, Bowman et al., 2014, Prior and Bowman, 2014, Searle and Chen, 2018, Toledo et al., 2011), temperature of the warmest quarter (Nicklen et al., 2019, Lebourgeois et al., 2014, Wright et al., 2018), isothermality (Zhang et al., 2013), and precipitation of the driest quarter (Toledo et al., 2011).

To account for the potential effect of biogeographic history, we associated plot data with the corresponding biogeographic regions. We adopted the biogeographic subdivision of the world proposed by Morrone (2015); unlike the historical biogeographic realms, which are based on animal species distribution, Morrone’s approach accounts for both plant and animal distribution and includes transition zones between the main biogeographic regions (henceforth ‘bioregions’; Morrone, 2015). The collated dataset included 12,557 plots, spanning across 48 countries, for a total of 1,056,431 tree observations and 8,206 species.

To minimize the influence of uneven plot density, we randomly selected one plot when multiple plots were found within one square kilometer of each other. In this way a total of 946 plots were removed. The final dataset included 11,400 plots and represented most of the world’s forested areas (Fig. 1a). It included the Australian (N = 1,962), Ethiopian (N = 40), Nearctic (N = 612), Neotropical (N = 2,561), Oriental (N = 127), and Palearctic (N = 2,753) bioregions, as well as the Mexican transition zone (N = 3,345; Fig. 1b).

**Figure 1.**
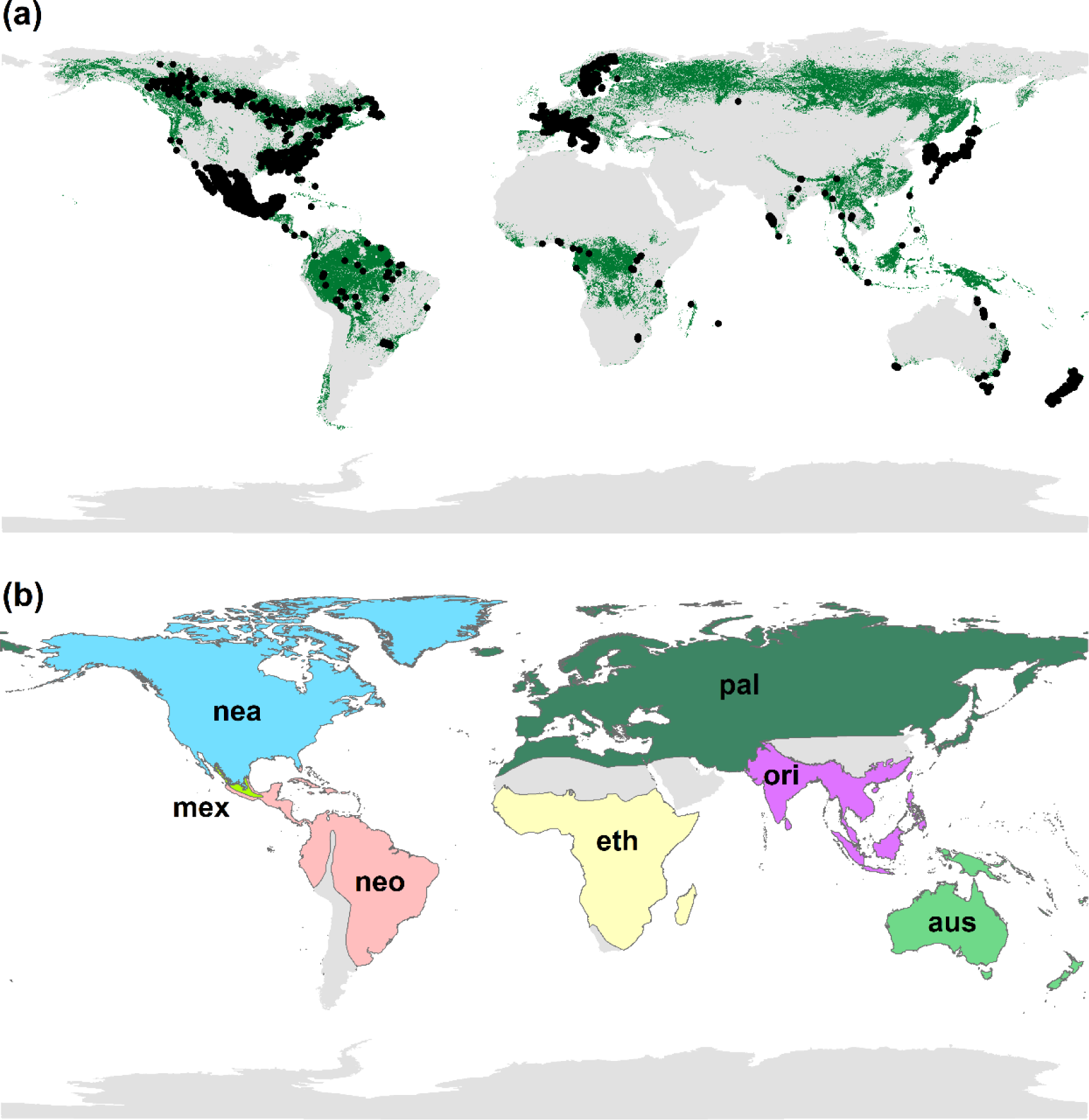
(a) Location of the forest plots used in this study (black dots) and global forest extent (green areas; from WWF forested areas 2017). (b) Bioregions included in this study (adapted from Morrone, 2015). Bioregion codes: aus: Australian bioregion, eth: Ethiopian bioregion, mex: Mexican transition zone, nea: Nearctic bioregion, neo: Neotropical bioregion, ori: Oriental bioregion, and pal: Palearctic bioregion.

### Statistical analyses

We used Generalized Additive Models (GAMs) to evaluate the relationship between forest structure and diversity on basal area. However, in such an analysis it is critical to account for unmeasured spatially structured effects as well as the influence of climate. To cope with the confounding between the spatial signal in the model residuals and that in the covariates, we applied the spatial+ approach introduced by Dupont et al. (2022), which has been shown to provide fixed effects estimates close to the true value in the case of spatial confounding (Urdangarin et al., 2023). In brief, spatial+ uses a two-stage smoothing spline model formulation to initially remove the spatial signal from the covariates before estimating their relationship to the response in the presence of residual spatial effects.

To prepare the data, the response variable (basal area), and the covariates species richness (Chao1), species evenness (Pielou’s J), and stem density were log transformed to stabilize the variance and to ensure positive definiteness when each variable is included as a response in the first modelling stage. To account for climate, we performed a Principal Component Analysis (PCA) using the available climate variables associated with each plot. We extracted axes 1 and 2 of the PCA (henceforth PCA1 and PCA2), which explained 43% and 29% of the variance respectively, and used them as covariates. For the first stage of regression, we fit a separate spatial model for each covariate, inclusive only of latitude and longitude as covariates. Given the broad distribution of the data across the Earth’s surface, we used isotropic smoothing splines on the sphere to model the spatial effects (Wood et al., 2016), setting the maximum basis dimension to 100. The residuals of these spatial models were then extracted and used in place of the original variables in the subsequent modelling step.

All candidate models in the second stage included as covariates species richness, species evenness, and stem density on basal area, and accounted for climate by including either annual values (mean annual temperature and total annual precipitation), quarterly extremes (mean temperature of warmest quarter and precipitation of wettest quarter) or the results of the PCA analysis, which combined all relevant climate variables (PCA1 and PCA2). To account for spatial effects in the response (basal area) we used smoothing splines on the sphere as described above for the first stage. A set of candidate models were designed to compare linear versus non-linear trends for the fixed effects, and to compare global versus regional effects for both the fixed effects and the residual variances. The latter specifies a location-scale GAM (Rigby and Stasinopoulos, 2005), also called distributional regression (Klein et al., 2015). Regional effects were specified using bioregions as a grouping parameter. The model set was characterized by the different use of the regional grouping as follows:

- Global linear model: a single slope for each covariate with no regional effects and a common variance.
- Global non-linear: a single smoothing spline for each covariate with no regional effects and a common variance.
- Regional linear model: random slopes for each covariate grouped by region and a common variance.
- Regional splines model: regional splines and a common variance
- Regional location-scale linear model: random slopes for each covariate and random intercepts for the variances, grouped by region.
- Regional location-scale spline model: regional splines for each covariate and random intercepts for the variances, grouped by region.

Three model sets were generated, one for each type of climate covariate (annual, quarterly, or PCA-based), for a total of 18 models. Model selection was based on comparisons of conditional Akaike Information Criterion (AIC) which estimates expected Kullback-Leibler divergence using the effective degrees of freedom adjusted for smoothing parameter uncertainty (Wood et al., 2016). Given the *a priori* potential for non-linear relationships, we could have omitted the linear models from the outset (as being implausible) and made inferences directly from the spline-based models. However, since all available methods to deal with spatial confounding are derived from linear assumptions, we included the linear models for comparison. The two-stage structure of the spatial+ model makes it particularly amenable to the use of non-linear trends in the final stage and our approach here closely resembles non-linear strategies used in the context of two-stage instrumental variables models (Marra and Radice, 2011, Briseño Sanchez et al., 2020). All analyses were done in R, using the packages ‘nlme’ (Pinheiro et al., 2023), ‘mgcv’ (Wood, 2019), and ‘tidyverse’ (Wickham et al., 2019). The code used to build and compare the models is provided in Supplementary Material 2.

## Results

We found substantial geographic differences in plot characteristics. Maximum plot size varied between bioregions, ranging from as small as 6 ha in the Nearctic to as large as 50 ha in the Neotropical bioregion. Despite this, the median and minimum values were similar across bioregions, except for the Oriental bioregion, for which median values were approximately one order or magnitude larger than the others (Table 1). Median basal area values were also similar, except for the Australian bioregion, for which substantially higher median values were obtained. Differences were detected in median species richness, which were lower in the Australian, Mexican, Nearctic, and Palearctic bioregions compared with the Ethiopian, Neotropical, and Oriental bioregions. Stem density also varied, with median values ranging between 347 stems/ha (Oriental bioregion) to 1,030 stems/ha (Neotropical bioregion; Table 1). Median, minimum, and maximum species evenness were comparable across bioregions.

**Table 1.**
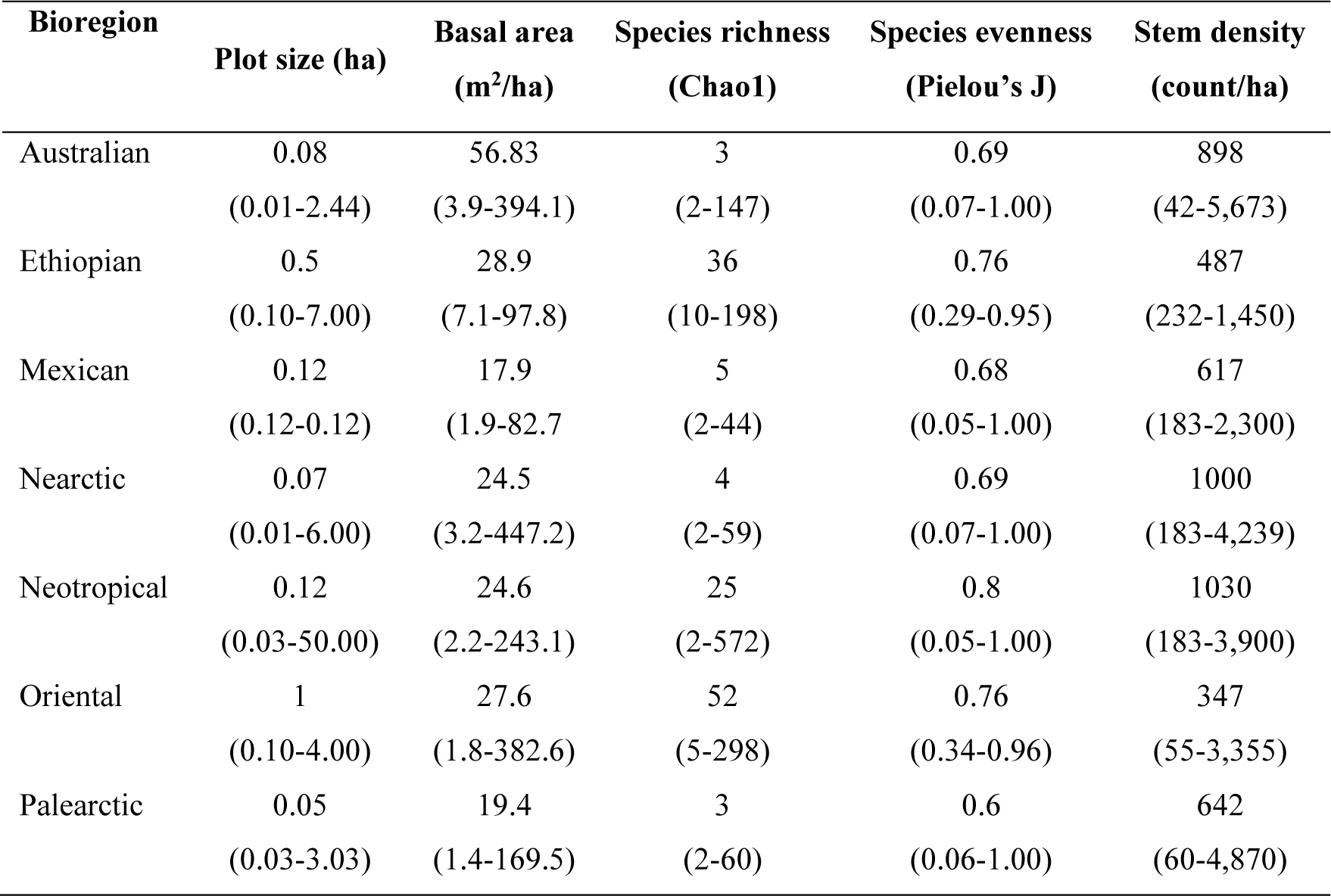
Summary statistics for each bioregion included in this study. Median values are presented, with the range reported between parentheses.

| Bioregion | Plot size (ha) | Basal area<br>(m<sup>2</sup>/ha) | Species richness<br>(Chao1) | Species evenness<br>(Pielou's J) | Stem density<br>(count/ha) |
| --- | --- | --- | --- | --- | --- |
| Australian | 0.08<br>(0.01-2.44) | 56.83<br>(3.9-394.1) | 3<br>(2-147) | 0.69<br>(0.07-1.00) | 898<br>(42-5,673) |
| Ethiopian | 0.5<br>(0.10-7.00) | 28.9<br>(7.1-97.8) | 36<br>(10-198) | 0.76<br>(0.29-0.95) | 487<br>(232-1,450) |
| Mexican | 0.12<br>(0.12-0.12) | 17.9<br>(1.9-82.7) | 5<br>(2-44) | 0.68<br>(0.05-1.00) | 617<br>(183-2,300) |
| Nearctic | 0.07<br>(0.01-6.00) | 24.5<br>(3.2-447.2) | 4<br>(2-59) | 0.69<br>(0.07-1.00) | 1000<br>(183-4,239) |
| Neotropical | 0.12<br>(0.03-50.00) | 24.6<br>(2.2-243.1) | 25<br>(2-572) | 0.8<br>(0.05-1.00) | 1030<br>(183-3,900) |
| Oriental | 1<br>(0.10-4.00) | 27.6<br>(1.8-382.6) | 52<br>(5-298) | 0.76<br>(0.34-0.96) | 347<br>(55-3,355) |
| Palearctic | 0.05<br>(0.03-3.03) | 19.4<br>(1.4-169.5) | 3<br>(2-60) | 0.6<br>(0.06-1.00) | 642<br>(60-4,870) |

### Model selection

Based on AIC values, all models using PCA1 and PCA2 as climate covariates consistently performed better than models using annual values or quarterly extremes, irrespective of model structure. The ‘Regional location-scale spline model’, which is the most complex of the models specifying non-linear regional effects for stem density, species richness, species evenness, and climate and included random intercepts by region for the scale, was selected as the best model (min delta AIC > 218) and explained 72% of the variation in basal area. We use the (selected) full model for all inferences which mitigates selection-induced bias that can otherwise arise when making inferences after model selection (Yates et al., 2023). Model summaries and model-selection results are presented in Supplementary Material 3. For a comparison, the linear effect of stand density, species richness, and species evenness is shown in Fig. S3.1 in Supplementary Material 3.

### Influence of forest structure and diversity on basal area

We found a clear relationship between stand density and basal area across all bioregions, with higher stand density consistently corresponding to higher values of basal area (Fig. 2).

**Figure 2.**
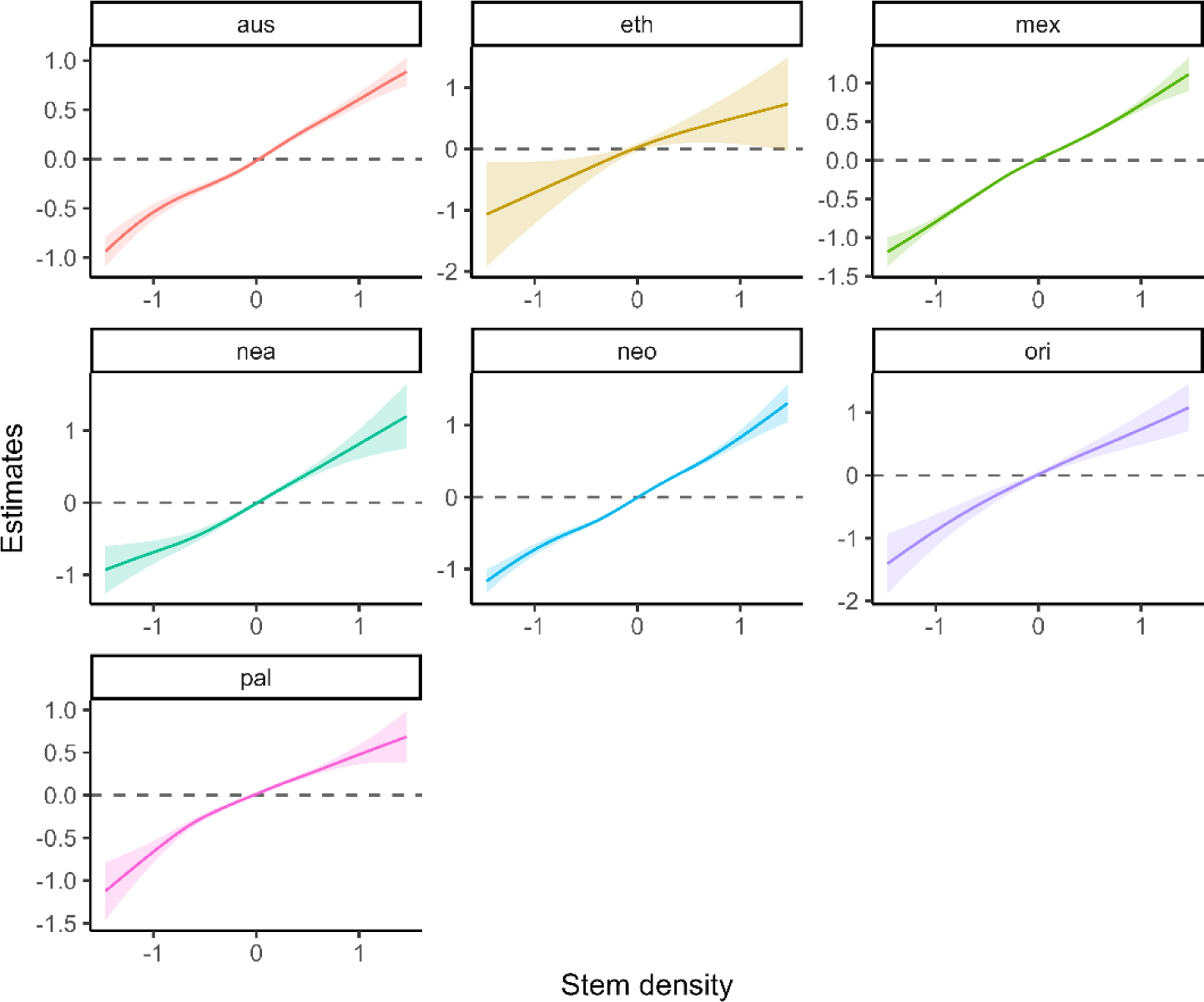
Smooth estimates of the partial effect of stem density on tree basal area across bioregions. Bioregion codes: aus: Australian bioregion, eth: Ethiopian bioregion, mex: Mexican transition zone, nea: Nearctic bioregion, neo: Neotropical bioregion, ori: Oriental bioregion, and pal: Palearctic bioregion.

Conversely, the effect of species richness on basal area varied across bioregions in both the strength and direction of the relationship. When accounting for uncertainty, four distinct trends could be identified: i) in the Palearctic, Nearctic, and Neotropical bioregions, basal area increased with species richness for lower basal area values, but with further increases in species richness not corresponding to additional increments in basal area (Fig. 3a); ii) in the Ethiopian and Oriental bioregions, species richness did not influence basal area (although their sample sizes were much smaller than in other regions); iii) in the Mexican transition zone, increases in species richness initially correspond to a decrease in basal area but subsequent increases in species richness have further effect, although that initial signal is characterized by high uncertainty; iv) the Australian bioregion was the only one where a negative relationship between species richness and basal area was detected (Fig. 3a).

**Figure 3.**
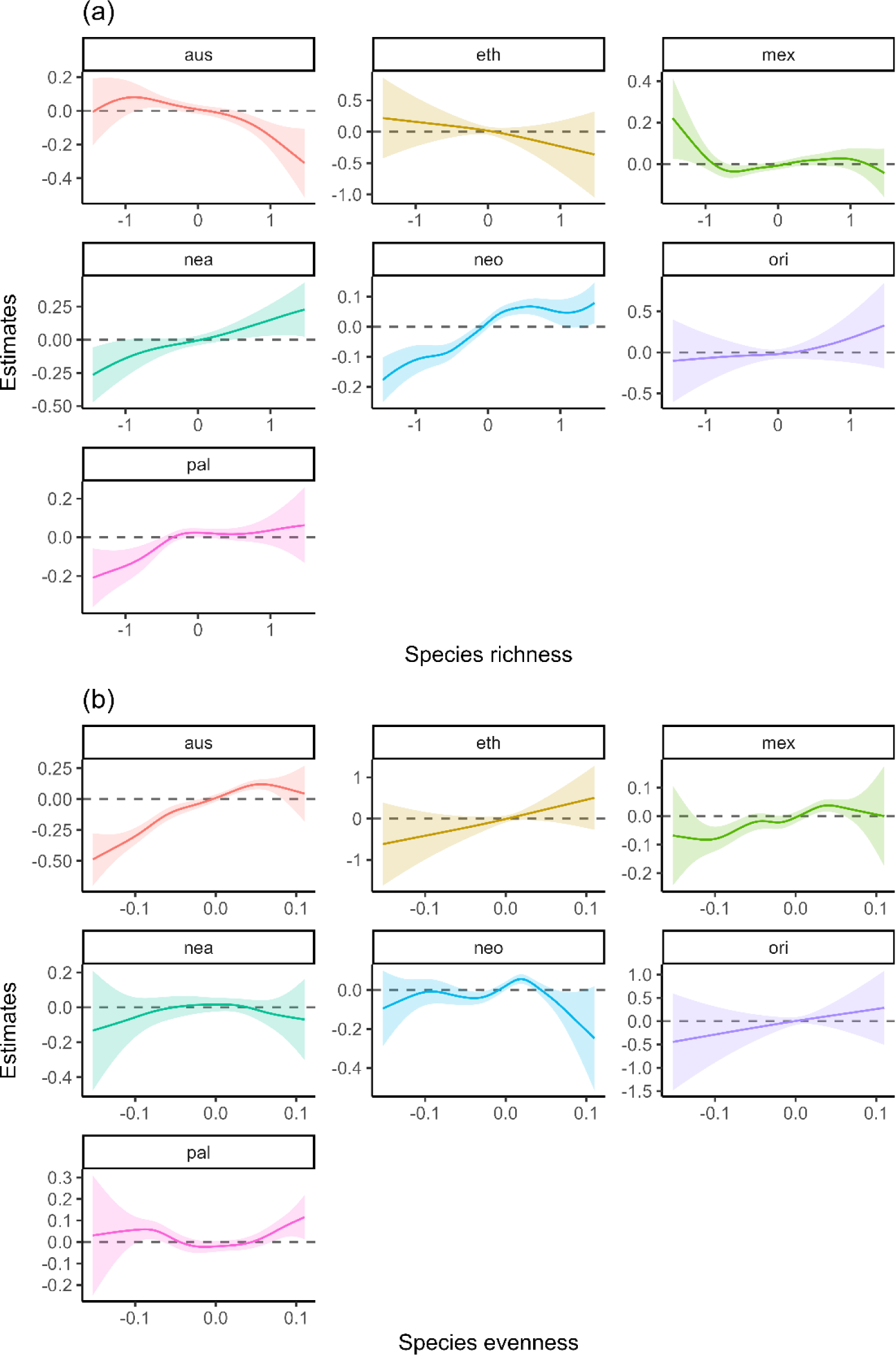
Smooth estimates of the partial effect of (a) species richness and (b) species evenness on tree basal area across bioregions. Bioregion codes: aus: Australian bioregion, eth: Ethiopian bioregion, mex: Mexican transition zone, nea: Nearctic bioregion, neo: Neotropical bioregion, ori: Oriental bioregion, and pal: Palearctic bioregion.

The effect of species evenness on basal area was moderate with only one geographic pattern; the Australian bioregion showed clear signal that more even forests were associated with higher basal area, while there was only a marginal increase in basal area was detected in the Ethiopian and Mexican bioregions and a slight decrease in the Neotropical bioregion (Fig. 3b). No relationship between these two variables could be identified in the Nearctic, Oriental, and Palearctic bioregions.

## Discussion

This study investigated the association between forest characteristics and basal area, which we used as a proxy for aboveground biomass. Our results supported the hypothesis that stem density, and to a certain extent species richness and evenness, are correlated with forest biomass. While the effects of stem density and species evenness were similar across bioregions, marked differences in the species richness-basal area associations were found across bioregions, suggesting that those dynamics are mediated by local drivers.

The strong and geographically consistent association found between stem density and basal area aligns with previous research (Ligot et al., 2018, Schietti et al., 2016, Sagar and Singh, 2006, Kara and Keleş, 2023), but it stands in contrast to the assumption that competition leads to self-thinning processes (Westoby, 1984, Ali et al., 2019). It is important to note that these processes unfold over time, and forest age has been found to influence biomass (Fang et al., 2014). Although we did not account for forest age in this study, as age data was not available for all plots, we consider it improbable that our results might be caused by a bias in the dataset towards younger forests. First, care was taken into selecting only non-disturbed sites and removing non-mature secondary forests. Further, the high number of plots included, as well as the variety of data sources, also makes it unlikely that an age bias was introduced. Plot size is also unlikely to have influenced this result, given that it was consistent across bioregions and characterized by very little model uncertainty (Fig. 2), despite substantial variation in plot size within and across bioregions (Table 1). Additionally, previous studies have detected no difference in the association between species richness and biomass across age classes (Caspersen and Pacala, 2001). A more straightforward and likely explanation is that stem density is strongly associated with basal area, despite the potential disproportionate influenced of large trees (Lutz et al., 2018).

Of the investigated forest characteristics, species richness was the one whose association with basal area showed the highest variability across bioregions. The overall positive association of species richness with basal area we found in the Palearctic, Nearctic, and Neotropical bioregion align with previous findings (Liang et al., 2007, Gamfeldt et al., 2013). An important caveat to that conclusion, however, is that the association was detected only for low values of basal area; as it increased, the correlated dissipated. Our results therefore only partially support the concept that the richness-biomass relationship is hump-shaped, and we could not find support for a decline in basal area at high-richness sites. The low association between species richness and basal area detected for bioregions whose extent is mostly limited to low latitudes is consistent with findings from previous studies, which showed that the strength of that relationships declines moving from high to lower latitudes (Wu et al., 2015). The negative trend detected for the Australian bioregion aligns with research showing that in Australian forests tree structural diversity (niche utilization) underpins biomass, unlike species diversity (Aponte et al., 2020). Notably, similarly contradictory results have been obtained for other habitats such as grasslands, where some studies have detected a positive relationship between species richness and biomass (Gross et al., 2014, Pastore et al., 2021), while others found no clear relationship between the two (Adler et al., 2011). This suggests that the richness-biomass association is likely mediated by other factors, such as environmental conditions, species characteristics, and biogeographic affinity (Noreika et al., 2019). This is particularly important as species richness is not only relevant for biomass accumulation, but it also support ecosystem services (Molina-Venegas et al., 2021), and is a critical conservation target to address the ongoing biodiversity crisis (Díaz et al., 2020).

Our results provided minor support to previous research concluding that evenness is positively associated with forest biomass (Zhang et al., 2012). This correlation was, however, detected only in a minority of bioregions and, when accounting for inferential uncertainty, the only clear signal was observed in the Australian bioregion. This is in contrast with research showing that in Australia lower evenness is associated with higher biomass, due to the high productivity of eucalypts, the dominant genus of most Australian forests (Gerwin et al., 2020). It does, however, agree with research from New Zealand, the other country included in the Australian bioregion, where Carswell et al. (2012) found a positive association between evenness and aboveground biomass, albeit in woody successions rather than established forests. Yet, an analysis of chrono-sequences across multiple countries found both positive and negative evenness-basal area associations in New Zealand depending on site location, and a positive effect in Australia (Wardle et al., 2008). Overall, this somewhat confusing pattern suggests that the effect of evenness on biomass is either i) negligible, which appears improbable given the substantial body of scientific literature which identified a relationship between these two variables (Zhang et al., 2012, Erskine et al., 2006, Lee et al., 2022), or, more likely, ii) it operates at more local scale than bioregional scale, as self-thinning processes depend strongly on forest type and species resource-use strategies (Urgoiti et al., 2023, Mulder et al., 2004, Gerwin et al., 2020, Rohr et al., 2016).

One limitation of this study is the small size of many of the plots included and the potential effect on biomass and diversity estimates. Studies have shown that accuracy of biomass estimates increased with plot size, although a strong relationship between the two parameters was found in plots as small as 0.05 ha (Kachamba et al., 2017). This suggests that the plots included in our analyses are likely sufficient to reliably estimate basal area (and thus biomass). The potential effect of plot size on species richness was addressed by using the widely used Chao1 richness estimator (Chao and Chiu, 2016) instead of species counts, as the latter has been shown to be influenced by plot size when establishing its association with biomass (Chisholm et al., 2013). It should be noted that, because of the global scope of this study, it was not feasible to identify exotic and invasive species within each dataset. It is thus possible that some plots included exotic species, which could lead to forest degradation (Asner et al., 2008) and have long-term effects on forest biomass that could not be detected in this study. Finally, although comprehensive, our analysis could not model some of the smaller bioregions outlined by Morrone (2015) because sufficient data could not be found for those areas (Andean and Cape bioregions, and Chinese and South American transition zones).

Future work could leverage on the growing body of long-term forest studies to evaluate temporal changes on the effect of forest structure and diversity on basal area, accounting for climate change and its effect on tree growth, as well as changing trends in disturbance regimes, such as the increased frequency and severity of wildfires, especially in areas not historically fire prone (Pearce, 2018, Bowman et al., 2020). Capitalizing on the increasing accuracy and availability of remotely sensed data can also complement field-based observations. For instance, LiDAR data can support assessment of vegetation volume and structure and thus support spatially explicit analyses of vegetation distribution (Camarretta et al., 2020), whilst high-resolution satellite imagery can provide information ongoing monitoring of vegetation spectral characteristics (Wallace et al., 2006). The influence of natural disturbance on forest characteristics should also be incorporated in future models. Revealing the finer-grained details of the influence of climate change, novel fire regimes on forest dynamics would be valuable to applied work that seeks to support biomass accumulation and protect biodiversity. Improving our understanding of the forest characteristics associated with tree biomass can support the design of effective management strategies for nature conservation and contribute to addressing the ongoing climate crisis.

## Acknowledgements

This study was funded by Australian Research Council Laureate Fellowship FL160100101 awarded to BWB. The authors would like to thank the managers of the Mexican, South Korean, Swedish, and Canadian forest inventories for supplying data not freely available online, and Mervyn Lötter for providing unpublished data for South Africa. We are also grateful to Cristian Montalvo-Mancheno and Sanghyun Hong for their help with translations in English from Spanish and South Korean.

## Author contributions

SO and LAY led study design. SO led data collection and preparation and wrote the manuscript, while LAY led data analysis, with assistance from RZA, and contributed to the writing. JCB, BWB, and JA conceived the initial concept, contributed to study design, and commented on the draft.

## Notes

### Competing Interest Statement

The authors have declared no competing interest.

