## Supplementary material 2 for "Tree species richness and evenness affect forest biomass differently across biogeographic regions": Supplementary material 2.R

```
#--------------------------------------------------------------------
#
# Global Forests 
#
# Apply the spatial+ method to the Global Forests model
#
# Dupont, E., Wood, S.N., and Augustin, N.H. (2022) Spatial+: 
# A novel approach to spatial confounding. Biometrics, 78: 1279– 1290. 
# https://doi.org/10.1111/biom.13656
#
# Author: L A Yates
# Date: 28t Feb 2023
# 
#--------------------------------------------------------------------

pkgs <- c("nlme","mgcv","tidyr","dplyr","readr","stringr","purrr", "ggbiplot") 
vapply(pkgs, library, logical(1), quietly = T, character.only = T, logical.return = T)

rm(list=ls(all=TRUE))

source("99_functions.R")

set.seed(8989)


#-----------
# Load data
#-----------

# load data
data_file <- "forests_thinned_2021_04_27.csv"
data_thinned <- read_csv(paste0("C:/Users/sondei/OneDrive - University of Tasmania/UTAS P Drive/Global Trees/Material from Luke Y/gf_fits_2023_03_06/",data_file))


#---------------------
# PCA
#---------------------

# Selected bioclimatic variables (from literature, see Methods)
bioSelected.pca <- prcomp(data_thinned[,c("bio_01", "bio_12", "bio_16", "bio_18", "bio_15", "bio_10", "bio_03", "bio_17")], 
                          center = TRUE, scale. = TRUE)   

bioSelected.plot <- ggbiplot(bioSelected.pca) # PCA1: 43.0% var. explained, PCA2: 29.1% var. explained

# Attach PCA values to dataset
bioSelected.PCA1_2 <- as.data.frame(bioSelected.pca$x[,1:2])
names(bioSelected.PCA1_2)[1:2] <- c("PCA1", "PCA2")
data_thinned <- cbind(data_thinned, bioSelected.PCA1_2)


#---------------------
# Prepare data
#---------------------

forests <- prepare_data(data_thinned)

# spline basis parameters
k_spatial <- 100
k_pred <- 15
group <- "reg" 
clim <- "PCA" # Change to 'annual' or 'quarter' to test the other climate variables

model_suffix <- paste0("models_",group, "_",clim,"_k.sp.",k_spatial,"_k.",k_pred)
message(paste("Model suffix:",model_suffix))

# List covariates
# Change 'PCA1' and 'PCA2' to:
#  -  'bio_01' and 'bio_12' to run models with mean annual temperature and total annual precipitation
#  -  'bio_10' and 'bio_16' to run models with tempaerature of the warmest quarter and precipitation of the wettest quarter
covars <- c("l_dens","l_chao","l_even","PCA1","PCA2")                 
message(paste("Model covariates:", paste(covars, collapse = ", ")))


#---------------------
# Stage-1 regressions
#---------------------

message("Fitting stage-1 models")

make_formula <- function(x,k){
  paste0(x, " ~ 1 + s(lat,lon, k = ", k_spatial,", bs = 'sos')") %>% 
  as.formula()
}

# fit stage-1 models for the confounded covariates
fits_stage_1 <- 
  covars %>% 
  {`names<-`(.,.)} %>% 
  map(make_formula, k = k_spatial) %>% 
  map(gam, data =  forests)
 
# extract residuals and add to data matrix 
forests_plus <- 
  fits_stage_1 %>% 
  {`names<-`(.,paste0("r_",names(.)))} %>% 
  map_dfc("residuals") %>% 
  bind_cols(forests) %>% 
  mutate(grp = forests[[group]])

#---------------------
# fit spatial+ models
#---------------------

message("Fitting global linear model")
# global effects, linear terms
m_glb_lin <- gam(l_ba ~ 
                  r_l_dens +
                  r_l_chao +
                  r_l_even +
                  r_PCA1 +
                  r_PCA2 +
                  s(lat,lon, k = k_spatial, bs = "sos"),
                data = forests_plus)

message("Fitting global spline model")
# global effects, non-linear terms
m_glb_spl <- gam(l_ba ~ 
                  s(r_l_dens, k = k_pred) +
                  s(r_l_chao, k = k_pred) +
                  s(r_l_even, k = k_pred) +
                  s(r_PCA1, k = k_pred) +
                  s(r_PCA2, k = k_pred) +
                  s(lat,lon, k = k_spatial, bs = "sos"),
                data = forests_plus)

message("Fitting group-level linear model")
# group effects, linear terms
m_grp_lin <- gam(l_ba ~ 0 +
                   grp +
                   r_l_dens:grp +
                   r_l_chao:grp +
                   r_l_even:grp +
                   r_PCA1:grp +
                   r_PCA2:grp +
                   s(lat,lon, k = k_spatial, bs = "sos"),
                 data = forests_plus)


message("Fitting group-level spline model")
# group effects, non-linear terms
m_grp_spl <- gam(l_ba ~ 
                   s(grp, bs = "re") +
                   s(r_l_dens, by = grp, id = 1, k = k_pred) +
                   s(r_l_chao, by = grp, id = 2, k = k_pred) +
                   s(r_l_even, by = grp, id = 3, k = k_pred) +
                   s(r_PCA1, by = grp, id = 4, k = k_pred) +
                   s(r_PCA2, by = grp, id = 5, k = k_pred) +
                   s(lat,lon, k = k_spatial, bs = "sos"),
                 data = forests_plus)


message("Fitting group-level location-scale linear model with random slopes")
# group effects, linear terms, location-scale model with random slopes
m_grp_lin_re_ls <- gam(list(l_ba ~ 1 +
                           s(grp, bs = "re") +
                           s(r_l_dens, grp, bs = "re") +
                           s(r_l_chao, grp, bs = "re") +
                           s(r_l_even, grp, bs = "re") +
                           s(r_PCA1, grp, bs = "re") +
                           s(r_PCA2, grp, bs = "re") +
                           s(lat,lon, k = k_spatial, bs = "sos"),
                         ~ 1 + s(grp, bs = "re")),
                    family = gaulss(),
                    data = forests_plus)

message("Fitting group-level location-scale spline model")
# group effects, non-linear terms, location-scale model (SLOW ~ 20 mins)
# (includes group-level variance estimates) 
m_grp_spl_ls <- gam(list(l_ba ~ 
                   s(grp, bs= "re") +      
                   s(r_l_dens, by = grp, id = 1, k = k_pred) +
                   s(r_l_chao, by = grp, id = 2, k = k_pred) +
                   s(r_l_even, by = grp, id = 3, k = k_pred) +
                   s(r_PCA1, by = grp, id = 4, k = k_pred) +
                   s(r_PCA2, by = grp, id = 5, k = k_pred) +
                   s(lat,lon, k = k_spatial, bs = "sos"),
                   ~ 1 + s(grp, bs = "re")),
                   family = gaulss(),
                   data = forests_plus)

message("Saving fits")
list(glb_lin = m_glb_lin,
     glb_spl = m_glb_spl,
     grp_lin = m_grp_lin,
     grp_spl = m_grp_spl,
     grp_lin_re_ls = m_grp_lin_re_ls,
     grp_spl_ls = m_grp_spl_ls) %>% 
  saveRDS(paste0("fits/", model_suffix,".rds"))

message("Done")


#---------------------
# Model comparison
#---------------------

# load models
models_PCA <- readRDS("fits/models_reg_PCA_k.sp.100_k.15.rds")
models_annual <- readRDS("fits/models_reg_annual_k.sp.100_k.15.rds")
models_quarter <- readRDS("fits/models_reg_quarter_k.sp.100_k.15.rds")

# model comparison
c(PCA = models_PCA, annual = models_annual, quarter = models_quarter) %>% 
  map_dbl(AIC) %>% {.-min(.)} %>% {.[order(.)]}
```
