## Supplementary material 3 for "Tree species richness and evenness affect forest biomass differently across biogeographic regions"

**Influence of forest structure and species diversity on tree basal area across biogeographic regions**

*Supplementary material 3*

- AIC results (Table S2.1)
- Linear effects (Fig. S2.1)
- Model summaries

**Table S3.1.** AIC values of the models including either i) mean annual temperature and total annual precipitation (prefix ‘annual’), ii) temperature of the warmest quarter and precipitation of the wettest quarter (prefix ‘quarter’), or iii) PCA axes 1 and 2 (prefix ‘PCA’) as climate covariates. See *Methods* section for a more detailed description of each model.

| **Model name** | **Model description** | **Climate covariate** | **AIC** |
| --- | --- | --- | --- |
| PCA.grp_spl_ls | Regional location-scale spline | PCA | 0 |
| PCA.grp_lin_re_ls | Regional location-scale linear | PCA | 218.7997 |
| PCA.grp_spl | Regional splines | PCA | 641.3461 |
| PCA.grp_lin | Regional linear | PCA | 975.6413 |
| PCA.glb_spl | Global non-linear | PCA | 1284.371 |
| PCA.glb_lin | Global linear | PCA | 1457.157 |
| quarter.grp_spl_ls | Regional location-scale spline | Quarter | 7088.875 |
| annual.grp_spl_ls | Regional location-scale spline | Annual | 7111.662 |
| quarter.grp_lin_re_ls | Regional location-scale linear | Quarter | 7361.984 |
| annual.grp_lin_re_ls | Regional location-scale linear | Annual | 7411.508 |
| quarter.grp_spl | Regional splines | Quarter | 7707.455 |
| annual.grp_spl | Regional splines | Annual | 7719.078 |
| quarter.grp_lin | Regional linear | Quarter | 8123.391 |
| annual.grp_lin | Regional linear | Annual | 8161.69 |
| quarter.glb_spl | Global non-linear | Quarter | 8380.59 |
| annual.glb_spl | Global non-linear | Annual | 8409.414 |
| annual.glb_lin | Global linear | Annual | 8600.807 |
| quarter.glb_lin | Global linear | Quarter | 8612.576 |

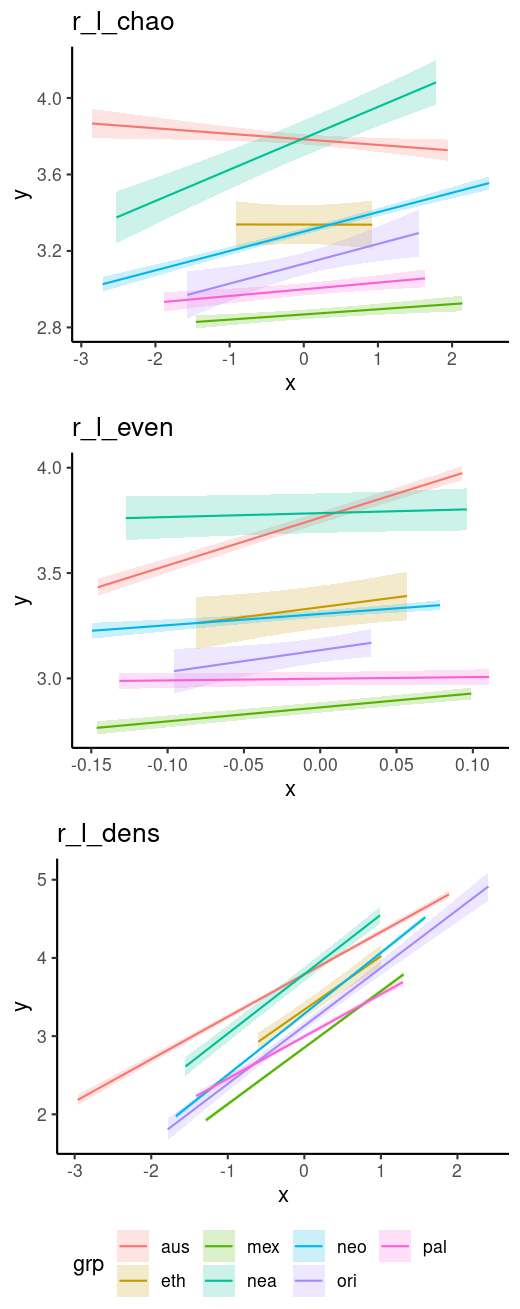
**Figure S3.1.** Linear effects of species richness (r_l_chao), species evenness (r_l_even), and stem density (r_l_dens) on forest basal area across different bioregions. Bioregion codes: aus: Australian bioregion, eth: Ethiopian bioregion, mex: Mexican transition zone, nea: Nearctic bioregion, neo: Neotropical bioregion, ori: Oriental bioregion, and pal: Palearctic bioregion.

**Model summaries. Summaries of the models included in the model set in which PCA axes were used as climate covariate.**

**Regional location-scale spline model**

```
Family: gaussian
Link function: identity

Parametric coefficients:
              Estimate Std. Error z value Pr(>|z|)
(Intercept)    3.14824    0.03612   87.17   <2e-16 ***
(Intercept).1 -0.93903    0.09256  -10.14   <2e-16 ***
---
Signif. codes:  0 ‘***’ 0.001 ‘**’ 0.01 ‘*’ 0.05 ‘.’ 0.1 ‘ ’ 1

Approximate significance of smooth terms:
                      edf Ref.df    Chi.sq  p-value
s(grp)              1.875  7.000    50.224  < 2e-16 ***
s(r_l_dens):grpaus  4.602  5.809   746.520  < 2e-16 ***
s(r_l_dens):grpeth  1.172  1.316    11.566 0.007578 **
s(r_l_dens):grpmex  4.906  6.189  1597.273  < 2e-16 ***
s(r_l_dens):grpnea  2.603  3.355   190.069  < 2e-16 ***
s(r_l_dens):grpneo  4.707  6.051  1368.506  < 2e-16 ***
s(r_l_dens):grpori  2.141  2.726    71.991  < 2e-16 ***
s(r_l_dens):grppal  3.642  4.665   432.999  < 2e-16 ***
s(r_l_chao):grpaus  3.825  4.937    22.589 0.000359 ***
s(r_l_chao):grpeth  1.177  1.327     0.985 0.350991
s(r_l_chao):grpmex  4.914  6.213    17.616 0.010240 *
s(r_l_chao):grpnea  3.419  4.374    14.887 0.006996 **
s(r_l_chao):grpneo  6.672  8.275   100.121  < 2e-16 ***
s(r_l_chao):grpori  1.489  1.847     0.870 0.521174
s(r_l_chao):grppal  3.894  4.967    15.718 0.008380 **
s(r_l_even):grpaus  3.880  4.889   126.003  < 2e-16 ***
s(r_l_even):grpeth  1.141  1.266     2.526 0.141995
s(r_l_even):grpmex  5.237  6.568    32.071 3.37e-05 ***
s(r_l_even):grpnea  2.904  3.682     1.659 0.763047
s(r_l_even):grpneo  4.441  5.620    33.760 5.36e-06 ***
s(r_l_even):grpori  1.156  1.291     1.016 0.409886
s(r_l_even):grppal  4.520  5.666    15.504 0.017776 *
s(r_PCA1):grpaus    5.071  6.461     8.045 0.287827
s(r_PCA1):grpeth    1.215  1.395     3.198 0.072224 .
s(r_PCA1):grpmex    5.219  6.554   490.645  < 2e-16 ***
s(r_PCA1):grpnea    3.104  3.974    31.544 2.42e-06 ***
s(r_PCA1):grpneo    6.185  7.759   185.205  < 2e-16 ***
s(r_PCA1):grpori    2.605  3.317     0.616 0.876671
s(r_PCA1):grppal    4.521  5.728   148.904  < 2e-16 ***
s(r_PCA2):grpaus    3.544  4.637     2.259 0.761486
s(r_PCA2):grpeth    1.140  1.259     0.228 0.850156
s(r_PCA2):grpmex    3.187  4.075     0.835 0.938272
s(r_PCA2):grpnea    1.265  1.486     1.584 0.261311
s(r_PCA2):grpneo    5.160  6.394    18.802 0.004672 **
s(r_PCA2):grpori    3.416  4.209     3.687 0.522278
s(r_PCA2):grppal    2.686  3.498    13.365 0.012499 *
s(lat,lon)         94.999 99.000 29954.442  < 2e-16 ***
s.1(grp)            5.740  6.000   809.563  < 2e-16 ***
---
Signif. codes:  0 ‘***’ 0.001 ‘**’ 0.01 ‘*’ 0.05 ‘.’ 0.1 ‘ ’ 1

Deviance explained = 72.1%
-REML = 5232.2  Scale est. = 1         n = 11400
```

**Global linear model**

```
Family: gaussian
Link function: identity

Formula:
l_ba ~ r_l_dens + r_l_chao + r_l_even + r_PCA1 + r_PCA2 + s(lat,
    lon, k = k_spatial, bs = "sos")

Parametric coefficients:
            Estimate Std. Error t value Pr(>|t|)
(Intercept) 3.165431   0.003706 854.113   <2e-16 ***
r_l_dens    0.639674   0.010338  61.876   <2e-16 ***
r_l_chao    0.076406   0.008699   8.783   <2e-16 ***
r_l_even    0.861412   0.085805  10.039   <2e-16 ***
r_PCA1      0.235457   0.008629  27.288   <2e-16 ***
r_PCA2      0.071080   0.006377  11.146   <2e-16 ***
---
Signif. codes:  0 ‘***’ 0.001 ‘**’ 0.01 ‘*’ 0.05 ‘.’ 0.1 ‘ ’ 1

Approximate significance of smooth terms:
             edf Ref.df     F p-value
s(lat,lon) 97.29     99 198.1  <2e-16 ***
---
Signif. codes:  0 ‘***’ 0.001 ‘**’ 0.01 ‘*’ 0.05 ‘.’ 0.1 ‘ ’ 1

R-sq.(adj) =  0.704   Deviance explained = 70.7%
GCV = 0.15801  Scale est. = 0.15658   n = 11400
```

**Global spline model**

```
Family: gaussian
Link function: identity

Formula:
l_ba ~ s(r_l_dens, k = k_pred) + s(r_l_chao, k = k_pred) + s(r_l_even,
    k = k_pred) + s(r_PCA1, k = k_pred) + s(r_PCA2, k = k_pred) +
    s(lat, lon, k = k_spatial, bs = "sos")

Parametric coefficients:
            Estimate Std. Error t value Pr(>|t|)
(Intercept) 3.165431   0.003672   862.1   <2e-16 ***
---
Signif. codes:  0 ‘***’ 0.001 ‘**’ 0.01 ‘*’ 0.05 ‘.’ 0.1 ‘ ’ 1

Approximate significance of smooth terms:
               edf Ref.df       F p-value
s(r_l_dens)  6.695  8.380 452.908  <2e-16 ***
s(r_l_chao) 10.604 12.338   9.155  <2e-16 ***
s(r_l_even)  7.075  8.748  12.689  <2e-16 ***
s(r_PCA1)    6.236  7.907 101.375  <2e-16 ***
s(r_PCA2)   13.117 13.850   7.407  <2e-16 ***
s(lat,lon)  97.188 99.000 185.486  <2e-16 ***
---
Signif. codes:  0 ‘***’ 0.001 ‘**’ 0.01 ‘*’ 0.05 ‘.’ 0.1 ‘ ’ 1

R-sq.(adj) =   0.71   Deviance explained = 71.3%
GCV = 0.15565  Scale est. = 0.15371   n = 11400
```

**Regional linear model**

```
Family: gaussian
Link function: identity

Formula:
l_ba ~ 0 + grp + r_l_dens:grp + r_l_chao:grp + r_l_even:grp +
    r_PCA1:grp + r_PCA2:grp + s(lat, lon, k = k_spatial, bs = "sos")

Parametric coefficients:
                  Estimate Std. Error t value Pr(>|t|)
grpaus           4.8188207  1.0293217   4.682 2.88e-06 ***
grpeth           1.2336883  1.0272847   1.201 0.229807
grpmex           2.9865796  0.7453330   4.007 6.19e-05 ***
grpnea           2.8890989  0.7453337   3.876 0.000107 ***
grpneo           2.7939734  0.7453705   3.748 0.000179 ***
grpori           3.3064917  1.0375281   3.187 0.001442 **
grppal           2.6201586  1.0440863   2.510 0.012103 *
grpaus:r_l_dens  0.5872370  0.0179217  32.767  < 2e-16 ***
grpeth:r_l_dens  0.6786979  0.1942089   3.495 0.000476 ***
grpmex:r_l_dens  0.7222756  0.0215801  33.469  < 2e-16 ***
grpnea:r_l_dens  0.7476142  0.0500378  14.941  < 2e-16 ***
grpneo:r_l_dens  0.7878046  0.0282144  27.922  < 2e-16 ***
grpori:r_l_dens  0.7670337  0.0646087  11.872  < 2e-16 ***
grppal:r_l_dens  0.5113917  0.0220585  23.183  < 2e-16 ***
grpaus:r_l_chao -0.0683528  0.0209493  -3.263 0.001107 **
grpeth:r_l_chao -0.1870491  0.1798380  -1.040 0.298317
grpmex:r_l_chao  0.0276644  0.0192257   1.439 0.150199
grpnea:r_l_chao  0.1910762  0.0391229   4.884 1.05e-06 ***
grpneo:r_l_chao  0.1073594  0.0162581   6.603 4.20e-11 ***
grpori:r_l_chao  0.1687143  0.0890278   1.895 0.058108 .
grppal:r_l_chao  0.0405454  0.0194995   2.079 0.037612 *
grpaus:r_l_even  2.3683888  0.1819684  13.015  < 2e-16 ***
grpeth:r_l_even  4.6742201  2.6570733   1.759 0.078577 .
grpmex:r_l_even  0.6326209  0.1551632   4.077 4.59e-05 ***
grpnea:r_l_even  0.0061995  0.3330446   0.019 0.985149
grpneo:r_l_even  0.4875261  0.2873507   1.697 0.089795 .
grpori:r_l_even  2.3085538  1.9202481   1.202 0.229305
grppal:r_l_even  0.0653707  0.1463350   0.447 0.655086
grpaus:r_PCA1    0.0622060  0.0290413   2.142 0.032216 *
grpeth:r_PCA1    0.4258518  0.2111043   2.017 0.043692 *
grpmex:r_PCA1    0.3189138  0.0173496  18.382  < 2e-16 ***
grpnea:r_PCA1    0.1764008  0.0448340   3.935 8.39e-05 ***
grpneo:r_PCA1    0.1765639  0.0212660   8.303  < 2e-16 ***
grpori:r_PCA1    0.0135449  0.0829846   0.163 0.870347
grppal:r_PCA1    0.2200910  0.0168857  13.034  < 2e-16 ***
grpaus:r_PCA2   -0.0092057  0.0221887  -0.415 0.678235
grpeth:r_PCA2    0.2766546  0.1832506   1.510 0.131147
grpmex:r_PCA2   -0.0009019  0.0163899  -0.055 0.956117
grpnea:r_PCA2    0.0198769  0.0591845   0.336 0.736993
grpneo:r_PCA2    0.0402538  0.0127405   3.160 0.001585 **
grpori:r_PCA2    0.0135074  0.0285621   0.473 0.636284
grppal:r_PCA2    0.0488570  0.0188562   2.591 0.009581 **
---
Signif. codes:  0 ‘***’ 0.001 ‘**’ 0.01 ‘*’ 0.05 ‘.’ 0.1 ‘ ’ 1

Approximate significance of smooth terms:
             edf Ref.df     F p-value
s(lat,lon) 96.63     99 79.35  <2e-16 ***
---
Signif. codes:  0 ‘***’ 0.001 ‘**’ 0.01 ‘*’ 0.05 ‘.’ 0.1 ‘ ’ 1

R-sq.(adj) =  0.717   Deviance explained = 98.6%
GCV = 0.15149  Scale est. = 0.14965   n = 11400
```

**Regional spline model**

```
Family: gaussian
Link function: identity

Formula:
l_ba ~ s(grp, bs = "re") + s(r_l_dens, by = grp, id = 1, k = k_pred) +
    s(r_l_chao, by = grp, id = 2, k = k_pred) + s(r_l_even, by = grp,
    id = 3, k = k_pred) + s(r_PCA1, by = grp, id = 4, k = k_pred) +
    s(r_PCA2, by = grp, id = 5, k = k_pred) + s(lat, lon, k = k_spatial,
    bs = "sos")

Parametric coefficients:
            Estimate Std. Error t value Pr(>|t|)
(Intercept)   3.0572     0.3906   7.827 5.46e-15 ***
---
Signif. codes:  0 ‘***’ 0.001 ‘**’ 0.01 ‘*’ 0.05 ‘.’ 0.1 ‘ ’ 1

Approximate significance of smooth terms:
                      edf Ref.df       F  p-value
s(grp)              4.350  7.000   3.404 1.75e-06 ***
s(r_l_dens):grpaus  7.547  9.185 119.376  < 2e-16 ***
s(r_l_dens):grpeth  1.552  1.874   8.350 0.003866 **
s(r_l_dens):grpmex  6.802  8.388 133.840  < 2e-16 ***
s(r_l_dens):grpnea  4.433  5.616  45.777  < 2e-16 ***
s(r_l_dens):grpneo  6.391  8.020  99.027  < 2e-16 ***
s(r_l_dens):grpori  3.862  4.781  29.915  < 2e-16 ***
s(r_l_dens):grppal  6.004  7.475  73.483  < 2e-16 ***
s(r_l_chao):grpaus  6.386  7.986   4.702 8.39e-06 ***
s(r_l_chao):grpeth  1.557  1.900   0.399 0.610726
s(r_l_chao):grpmex  6.561  8.119   2.087 0.032779 *
s(r_l_chao):grpnea  5.327  6.640   2.894 0.005815 **
s(r_l_chao):grpneo  8.301 10.068   5.892  < 2e-16 ***
s(r_l_chao):grpori  2.680  3.392   2.271 0.109617
s(r_l_chao):grppal  6.104  7.587   3.304 0.001480 **
s(r_l_even):grpaus  5.797  7.214  26.345  < 2e-16 ***
s(r_l_even):grpeth  1.384  1.654   2.779 0.089330 .
s(r_l_even):grpmex  6.443  8.002   3.006 0.002269 **
s(r_l_even):grpnea  4.208  5.295   0.587 0.712226
s(r_l_even):grpneo  5.253  6.586   3.574 0.001036 **
s(r_l_even):grpori  1.622  2.009   0.077 0.927267
s(r_l_even):grppal  6.400  7.924   2.596 0.008132 **
s(r_PCA1):grpaus    8.855 10.664   2.101 0.020394 *
s(r_PCA1):grpeth    1.915  2.364   1.856 0.162535
s(r_PCA1):grpmex    7.809  9.412  36.337  < 2e-16 ***
s(r_PCA1):grpnea    5.648  6.978   7.069  < 2e-16 ***
s(r_PCA1):grpneo    8.842 10.666  10.954  < 2e-16 ***
s(r_PCA1):grpori    4.949  6.077   1.892 0.077612 .
s(r_PCA1):grppal    7.850  9.454  19.794  < 2e-16 ***
s(r_PCA2):grpaus   12.612 13.207   1.672 0.059607 .
s(r_PCA2):grpeth    4.251  4.858   0.627 0.702216
s(r_PCA2):grpmex    9.288  9.878   0.835 0.608923
s(r_PCA2):grpnea    6.536  7.336   6.305  < 2e-16 ***
s(r_PCA2):grpneo   11.872 12.269   1.699 0.058267 .
s(r_PCA2):grpori   11.794 12.801   2.083 0.010435 *
s(r_PCA2):grppal   10.045 10.720   3.515 0.000114 ***
s(lat,lon)         96.550 99.000  74.868 0.010673 *
---
Signif. codes:  0 ‘***’ 0.001 ‘**’ 0.01 ‘*’ 0.05 ‘.’ 0.1 ‘ ’ 1

R-sq.(adj) =   0.73   Deviance explained = 73.7%
GCV = 0.14721  Scale est. = 0.14309   n = 11400
```

**Regional location-scale linear**

```
Family: gaulss
Link function: identity logb

Formula:
l_ba ~ 1 + s(grp, bs = "re") + s(r_l_dens, grp, bs = "re") +
    s(r_l_chao, grp, bs = "re") + s(r_l_even, grp, bs = "re") +
    s(r_PCA1, grp, bs = "re") + s(r_PCA2, grp, bs = "re") + s(lat,
    lon, k = k_spatial, bs = "sos")
~1 + s(grp, bs = "re")

Parametric coefficients:
              Estimate Std. Error z value Pr(>|z|)
(Intercept)    3.15632    0.04632  68.143   <2e-16 ***
(Intercept).1 -0.90379    0.09300  -9.718   <2e-16 ***
---
Signif. codes:  0 ‘***’ 0.001 ‘**’ 0.01 ‘*’ 0.05 ‘.’ 0.1 ‘ ’ 1

Approximate significance of smooth terms:
                   edf Ref.df    Chi.sq p-value
s(grp)           2.035      6    2208.4  <2e-16 ***
s(r_l_dens,grp)  6.906      7  184286.9  <2e-16 ***
s(r_l_chao,grp)  5.342      7   26764.2  <2e-16 ***
s(r_l_even,grp)  5.070      7    2710.5  <2e-16 ***
s(r_PCA1,grp)    6.346      7   34091.0  <2e-16 ***
s(r_PCA2,grp)    3.386      7     321.7   0.163
s(lat,lon)      95.160     99 2042627.3  <2e-16 ***
s.1(grp)         5.751      6     792.6  <2e-16 ***
---
Signif. codes:  0 ‘***’ 0.001 ‘**’ 0.01 ‘*’ 0.05 ‘.’ 0.1 ‘ ’ 1

Deviance explained = 71.1%
-REML = 5283.3  Scale est. = 1         n = 11400
```
